# Global dissection of the impact of Alzheimer’s disease on brain architecture and behavior: High resolution MRH resolves robust regional effects

**DOI:** 10.1101/2025.01.08.632011

**Authors:** G. Allan Johnson, Yuqi Tian, Kathryn Hornburg, James J. Cook, Yi Qi, Leonard E. White, John T. Killmar, Catherine Kaczorowski, David G. Ashbrook, Robert W. Williams

**Author notes:** Correspondence to G. Allan Johnson. These authors contributed equally to this work (G. Allan Johnson, Yuqi Tian, Kathryn Hornburg).

## Abstract

Alzheimer’s disease (AD) affects brain regions with remarkable heterogeneity, but the precise impact of this disease on hundreds of small cortical, subcortical, and brainstem regions remains poorly defined. Here, as a prelude to testing preclinical models to prevent AD, we systematically quantified effects of human AD mutations in *APP* and *PSEN1* on 231 regions and comprehensively evaluated changes in volume with unprecedented resolution in genetically diverse mice as a function of sex and genetic background. We studied 34 5XFAD F1 hybrids and 23 sibling controls at 14 months, evaluating learning and memory behaviors, followed by *ex vivo* diffusion tensor images (DTI) at 25 µm resolution. We delineated 231 regions of interest (ROIs) bilaterally with high precision. Remarkably, we found bidirectional changes: marked volume increases (up to 10%) in neocortex, hippocampus, amygdala, and sensory nuclei, contrasted with decreases in striatum, pallidum, thalamus, hypothalamus, and most fiber tracts. These opposing effects are unrelated to amyloid load and are likely to reflect temporal gradients in susceptibility of ROIs. Effects are similar in both sexes but far more prominent in females. Genetic background strongly modulates penetrance of the human mutations with the AD-BXD77 F1 type having the greatest sensitivity. Light sheet microscopy and stereological analysis of NeuN+ neurons and amyloid β aggregates in 22 regions revealed up to 20% loss of cells in CA3 and in anterior and intralaminar parts of the thalamus. Volumetric changes correlated with impaired fear acquisition and memory, with cases and controls often showing opposite relations between performance and regional volumes. These findings reveal unprecedented regional heterogeneity in AD progression and suggest therapeutic efficacy may vary substantially across genetic backgrounds and between sexes.

## INTRODUCTION

Diagnostic criteria of Alzheimer’s disease (AD) are heterogeneous and controversial. This complexity arises from complex relations between cognitive and behavioral symptoms, neuroimaging, and neuropathological criteria^3,4,5^. The same is true of well-characterized mouse models of AD, in which there is considerable disagreement about major consequences of human familial autosomal dominant AD mutations on specific types of structural and functional outcomes^6–8^. One important factor emphasized by the work of both Murdy and colleagues and Gurdon and colleagues is that genetic backgrounds of AD mutation vary greatly and in ways that can make replication difficult^9,10^. Another problem is that there is limited systematic quantitative case-control data on effects of AD in either humans or mouse models—particularly on the integrity of cortical and subcortical regions as a function of mutation types, sex, age of onset of behavioral symptoms, environment, and treatment history. We do know that the strong role of genetic background effects and sex are key cofactors that must be incorporated into pre-clinical AD mouse models^11,12,13^. One of the best examples of the vital importance of genetic background and of sex in humans is the major differences in AD risk associated with APOE ε4 allele dosage^14,15,16^. In Asian ancestries this allele has strong deleterious effects with high odds ratios, but those with African and Hispanic ancestries have much less risk, even those with two copies of the ε4 variant—odds ratios as low as 1.26^15,17^. There is also a sex-by-genotype interaction effect of ε3/ε4 heterozygotes across several genetic ancestries with consistently higher risk for females^17^.

As in humans, both sex and genetic ancestry (strain genetic background) also have very strong effects and can even reverse directions of effects of mutations on behavior^18^. Reluctance to accept this complexity is a cause of many problems and misunderstandings. For example, in a recent MODEL-AD study of the 5XFAD mutation^19^ the authors recommend “the use of 5XFAD mice for AD studies focused on sex-based differences, immune regulation, and Aβ deposition *but not for translational studies evaluating the potential of therapeutics to treat cognitive impairments*” (emphasis added). This statement *must* be qualified by the important modifier: “on the background of the C57BL/6J inbred strain.” We have known since the work of Kaczorowski and colleagues^12 20 21^ that cognitive effects of the bigenic 5XFAD transgene (*APP* K670N, M671L, and I716V mutations; *PSEN1* M146L and L286V mutations) depends on strain background^22^. For example, Neuner and colleagues discovered 3-fold variation in contextual fear memory in AD-BXD F1s that is dependent on strain background. While AD-BXD81 F1s are impaired in a contextual fear memory even at 6 months of age, the AD-C57BL/6J mice are *the least affected of 26 types*. The contrast is even greater when quantifying differences between cases and controls at 14 months. Memory function degrades by 20–45% on 10 backgrounds but remains stable on four other backgrounds of these AD-BXD hybrids.

Here we extend work by Kaczorowski and colleagues^12^, the MODEL-AD consortium^23,24^, and the recent cytological and gene expression analyses of Gurdon and colleagues ^10^. We have systematically quantified the impact of the 5XFAD transgene on brain structure in both sexes using a case-control design at 14 months of age. We estimated changes in volumes of 231 cortical and subcortical regions and tracts bilaterally in 57 AD-BXD mice, both cases and sex-matched controls, using diffusion tensor imaging at 25 µm resolution—several hundred-fold higher than in previous work^24,19^. Our main goal is to expose the role of the human *APP* and *PSEN1* mutations in a tightly controlled laboratory setting, while evaluating their impact as a function of genetic background^13^ and sex.

Our second long-term goal is to set the stage for interventional studies to explore the preclinical efficacy of drugs to treat AD as early as possible, similar to the early interventions now being tested in human clinical cohorts^25,5^. To do this in a translationally robust way, we need to know much more about those regions that are affected first and most severely by AD. We also need to know which of hundreds of brain regions contribute to different types of cognitive loss and behavioral changes. Here we address four specific questions:

1. Can we create quantitative and comprehensive maps of the impact of the 5XFAD transgene on volumes of both large cortical and very small subcortical CNS regions and fiber tracts?
2. Can we demonstrate significant effects of genetic background and of sex on modulating the impact of the 5XFAD human mutations on regional volumes?
3. Can we demonstrate cellular correlates of changes in volume, in particular ab load and NeuN+ neuron densities and numbers?
4. Can we link volumetric changes in specific ROIs to key behavioral outcomes, particularly learning and memory?

## RESULTS

### Overview

We have used an analytic workflow similar in sequence to clinical and pathological investigations of humans with AD (Fig. 1; Ref ^26^). We start with behavioral assessments of general function and cognition and progress to MRI scans to confirm and refine diagnoses. In this workflow, we add postmortem analyses and evaluate changes at the cellular level^27–29^). However, unlike conventional studies of AD using mouse models, we do not rely on a single fully inbred strain, but instead incorporate a high level of genetic variation that matches that of human populations^30,31,32,12,22,20,33,10^. The AD-BXD F1s are replicable hybrids that can be used for well-balanced case-control designs to study the efficacy of pre-clinical treatments.

**Fig. 1.**
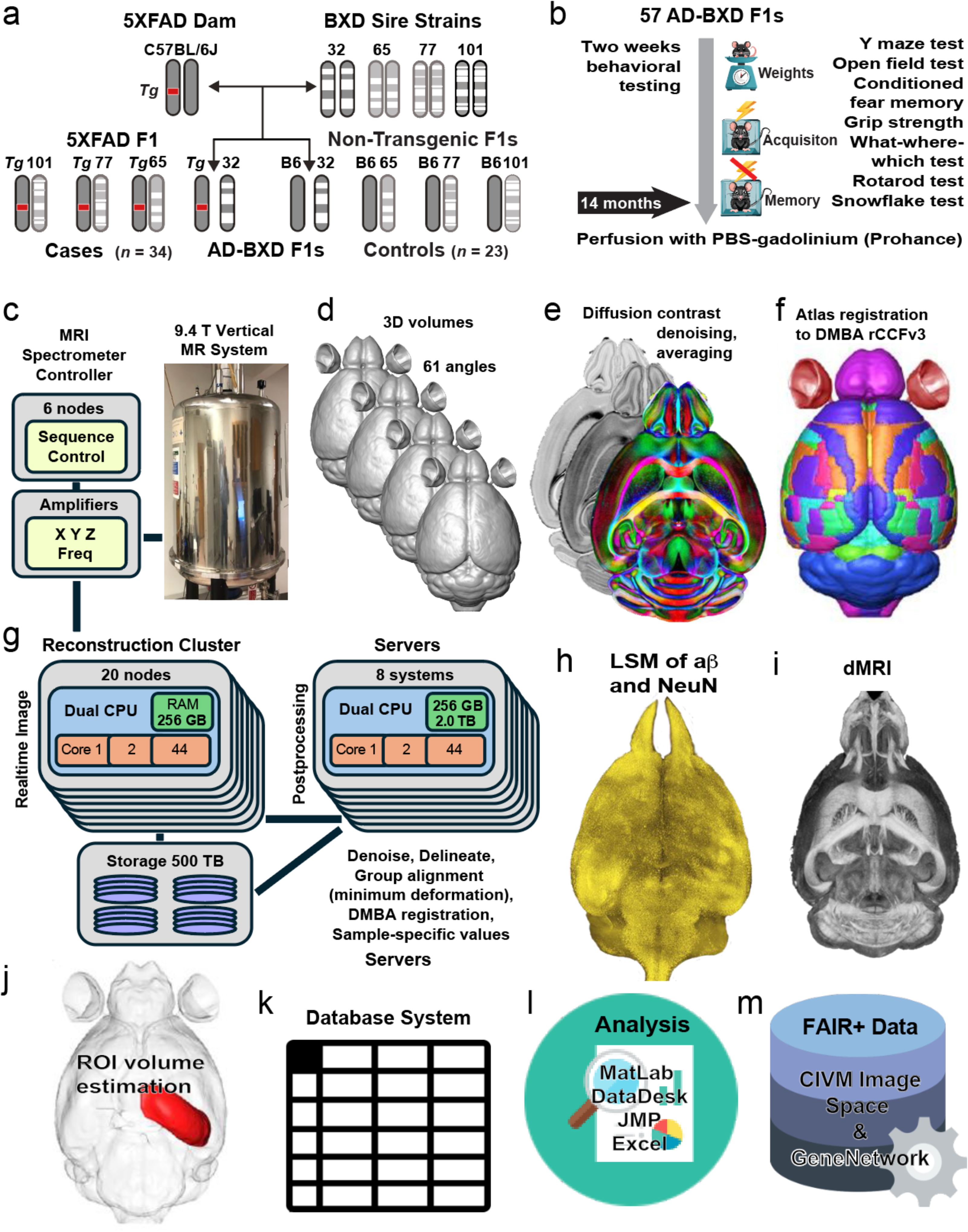
Workflow to evaluate impact of 5XFAD transgene effects on volumes of 231 brain regions. **a,** Female C57BL/6J mice hemizygous for the dominant 5XFAD transgene (Tg) were bred to males from four different BXD strains to generate genetically diverse, but isogenic F1 offspring—both Tg carriers (cases) and non-transgenic (NTg) controls. F1 offspring from a cross are genetically identical except for the presence of a transgene (red block). b, Behavioral and physiological phenotyping was carried out at 14 months of age. Contextual fear memory acquisition (CFA) was measured, and the contextual fear memory retention (CFM) was tested 24 hours later. Animals are perfusion fixed for MR histology and sent to the CIVM at Duke. **c,** Intact whole crania were scanned on a 9.4T vertical bore magnet using DTI with compressed sensing. A script controls the scanner and transfers the raw data to the cluster for reconstruction. **d,** 3D MRIs are generated at 25-µm resolution and at 61 angles (5 interleaved control scans), each with a different gradient. **e,** an integrated 4D volume of all scans is created, denoised, and transferred to DSI Studio. This 4D data set is used to generate diffusion tensor output types, including eight 3D image modalities generated from the DTI, including axial diffusivity (AD), mean diffusivity (MD), radial diffusivity (RA), fractional anisotropy (FA). **f,** The set of images are passed to the ANTs pipeline to register all volumes into the Duke Mouse Brain Atlas (DMBA) framework using a reduced subset of the Common Coordinate Framework (rCCFv3) labels. Once registration is complete, full volume delineations from the DMBA rCCFv3 are applied for all ROIs. The transformations with all delineations are then back-projected (inverted) to the original specimen image. We extract quantitative volumetric and other data types bilaterally from each ROI (absolute and normalized volume, MD, FA, etc.). **i,** Volumes with labels are passed back to DSI Studio and ROIs are seeded for tractography to generate connectomes. **j,** The brain is removed from the skull, cleared, stained for NeuN and ab and scanned with selective plane illumination. The LSM data sets (∼1TB total) are mapped into the undistorted MRI volume. Cell counts and ab load for selected ROIs are generated as in Tian et al. 2023. k, All data and metadata are stored in an Oracle database. **l,** Data sets are exported to MATLAB, JMP, CSV, R, or Excel for downstream analyses. **l,** We provide FAIR+ data access for full 3D images from the CIVM ImageSpace web service (https://civmimagespace.civm.duhs.duke.edu). All summarized ROI-level data are aggregated along with metadata, behavioral and morphometric data in *GeneNetwork.org* (select **Species** = Mouse; **Group** = BXD NIA Alzheimer’s Studies; and choose a data **Type**).

Results are presented for three major data types: 1. MRH: volume differences for 230 brain regions that we have delineated bilaterally, and of course, the whole brain, for 57 cases and controls; 2. light sheet microscopy: Corroborative whole brain stereology of the cytological impact of 5XFAD at 14 months in AD-BXD77 F1 cases (*n* = 9) and sibling controls (*n* of 7); and 3. behavioral analyses to probe the relations between learning and memory function with volumetric differences as a function of transgene status, genetic backgrounds, and sex.

### 1. Volumetric Analysis of the Impact of 5XFAD by MRH

#### The critical importance of accurate measurements of volume phenotypes

MRI of postmortem tissues at microscopic resolution of 25 µm—what we refer to as magnetic resonance histology^34^—can produce images at volumetric resolutions more than 1000-fold higher than that of *in vivo* studies^29^. The difference is striking. Figure 2a is from an optimized *in vivo* study by Crombé^1^ whereas Figure 2b is the much-improved *ex vivo* resolution we achieve using the current methods (**Fig. 1c–j**). Four major upgrades have made this possible:

1. Spatial resolution that far better than previous work using mouse models (**Fig. 2a,b**)
2. High-angular resolution diffusion imaging, 61 angles in our case
3. Multispectral segmentation of brain regions and tracts using DWI and fractional anisotropy (FA)
4. Whole brain 3D coverage using a set of 231 ROIs, that are consistent with the CCFv3 (Ref ^35^), but with stereotaxic correction^2^ (**Fig. 1f**)

**Fig. 2.**
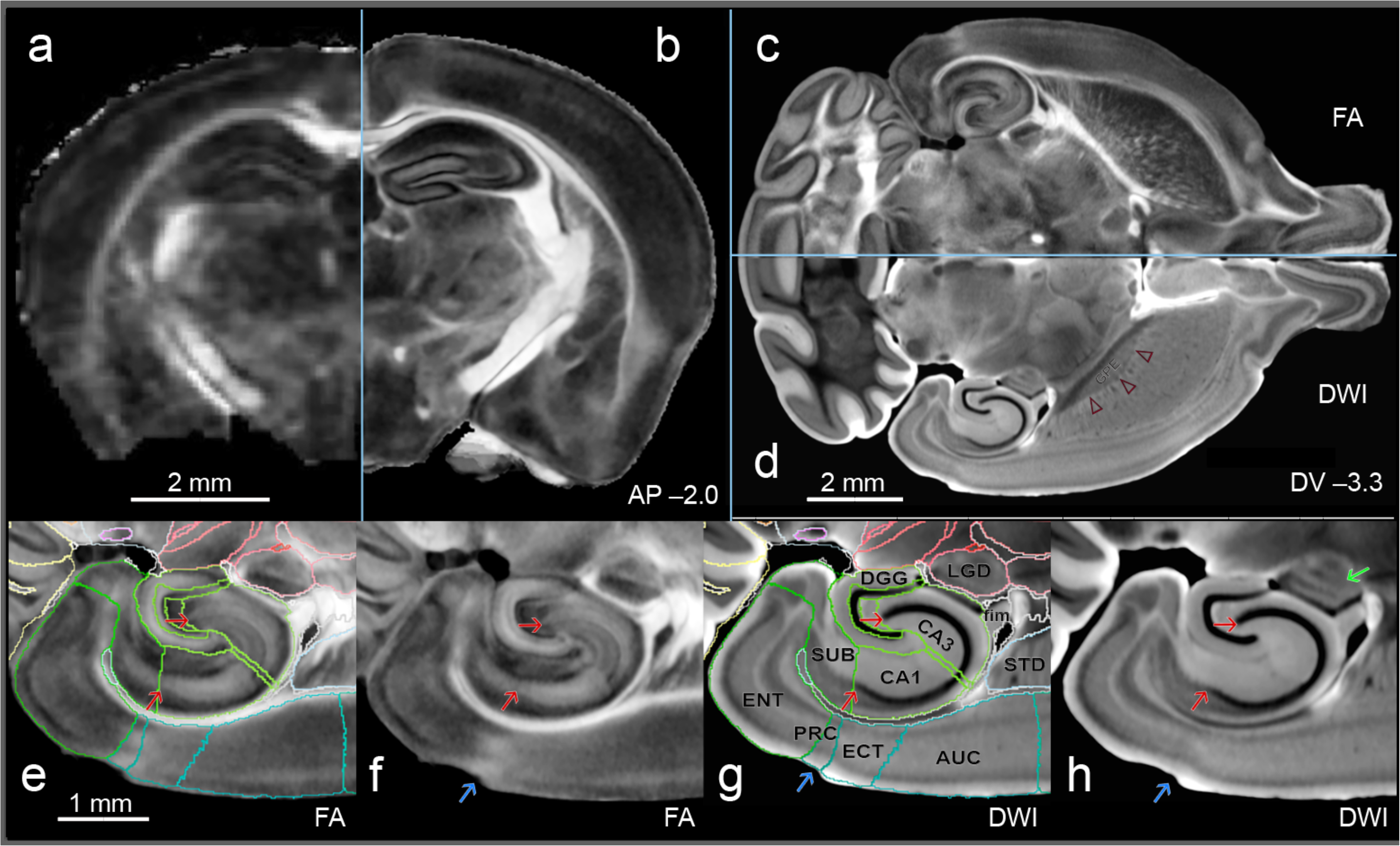
Spatial and contrast resolution increase precision in structure delineation. **a,b,** Side-by-side comparison of optimized fractional anisotropy (FA) *in vivo* MRI (left) and *ex vivo* MRH used in the present work (right) to emphasize the improvements in spatial resolution. Figure 2a is of a 5XFAD B6 congenic mouse from Ref^1^ at a voxel dimensions of 0.082 x 0.082 x 0.203 mm and voxel volume of 0.0014 mm^3^. **b,** the right side is a closely matched level of our current *ex vivo* protocol of an AD-BXD77 F1 specimen at an isotropic voxel volume of 25 µm^3^ (0.000016 mm^3^)—an 87-fold improvement in resolution. **c,d,** FA and DWI images are used in the present study to maximize the *contrast resolution* for the delineation of structural boundaries. Minimum deformation templates are computed for each individual as a function of its own AD-BXD F1 genetic background—four in this study. Images are registered to the FA and DWI consensus volume of the Duke Mouse Brain Atlas ^2^as shown in **d-h** for the minimum deformation template (average) of 7 AD-BXDF1 controls. Fiber tracts and subtle lamination differences in cortical and hippocampal layers are prominent in the FA images, but boundaries such as those between dorsal striatum (STD) and globus pallidus externa (GPE) are much more prominent in **d**—the DWI contrast (three outlined red arrowheads). **e-h,** Higher magnification pairs of FA and DWI cross-sections centered on the hippocampal formation. Sets of red arrows mark the start and end of the pyramidal cell layer extending from CA1 to CA3 and into the hilus of the dentate gyrus (labels in **g**). The final delineations combine information from these and other modalities. Three blue arrows in **f**, **g**, and **h** mark the caudal-lateral end of the perirhinal cortex (PRC). This boundary is well defined in by FA in **g**, but not by DWI in **h**. In **h** the green arrow points at a lamination pattern in the LGD that is rarely seen in any other dMRI modality. Other abbreviations: LGD: Lateral geniculate nucleus, dorsal; fim: fimbria; STD: Striatum, dorsal; AUC: Auditory cortex; ECT: Ectorhinal cortex; ENT: Entorhinal cortex; SUB: subiculum; CA1 and CA3 regions of hippocampus; DGG: Dentate gyrus. DMBA anterior-posterior (AP) and dorso-ventral (DV) levels of these sections are given in **b** and **d**.

#### No effects of 5XFAD mutation on total brain volume on any of four AD-BXD genetic backgrounds

We imaged 57 cases and controls belonging to four types of crosses of 5XFAD-by-BXD F1s, what we refer to as AD-BXD F1s, both the cases and their controls. Numbers of males and females are reasonably well-matched, 26 versus 31. However, the distribution is uneven among the AD-BXD F1 subtypes. The main consequence of the unevenness is variation in statistical power when analyses are stratified by genetic background. We intentionally included 50% more 5XFAD cases than controls (34 vs 23) because the transgene does increase within-class variance—neuroanatomical and behavioral.

and sex. Brain volumes of 5XFAD cases and their controls are 475.2 ± 3.45 mm^3^ SEM and 478.6 ± 6.14 mm^3^ SEM, respectively. Given that females are more affected by FAD in both humans and mice, we controlled for or stratified most statistical tests by sex. There is a –9.4 mm^3^ absolute difference between female cases and controls and a +1.4 mm^3^ difference between males. Neither difference is significant. Similarly, volumes of 26 males and 31 females are 472.9 ± 4.0 mm^3^ and 479.6 ± 4.8 mm^3^; again, not significant. Finally, even when stratified by the four F1 types (AD-BXD32 F1, AD-BXD65 F1, AD-BXD77 F1, and AD-BXD101 F1), there are still no significant differences in brain volume within type. Our conclusion is that here is no evidence of a global effect of the 5XFAD transgene on brain volume in AD-BXD mice at 14 months. In contrast, there is a strong genetic effect of strain background on overall brain volume. The AD-BXD32 F1s have the largest brain volumes (495 ± 4.9 mm^3^) whereas the AD-BXD101 F1s have the smallest (447 ± 4.1 mm^3^). However, there is again no significant effect of the 5XFAD transgene within any of the four AD-BXD types.

#### Strong effects of the 5XFAD transgene on regional volumes on all AD-BXD backgrounds

We compute CVs of independently segmented right and left ROIs—the standard deviation of the values divided by their mean—the y axis on figure 3c. The average CV of left-right comparisons of all 231 ROIs, including the two halves of the whole brain (BRN) is 1.8% (Fig. 3c). As expected, larger ROIs such as entire hemispheres, can be delineated more accurately, and consequently the correlation between volumes of ROIs and their CVs is –0.7—and this correlation is almost certainly due to technical error rather than true asymmetry. The minimum CV is 0.32% for brain hemispheres (BRN). The highest CVs are 5.7% for ROIs for very small nuclei such as the trapezoid body (NTB) and the paraventricular nucleus of the hypothalamus (PVH). These two ROIs have bilateral summed volumes of 0.084 mm^3^ and 0.25 mm^3^, respectively (**Fig. 3c**). However, at a volume of merely 1.0 mm^3^ or 1 mg of tissue, the CV of ROIs average about 2%, equivalent to 20 µg or 20 nanoliters of tissue. This level of replicability of right and left delineations gives us unprecedented accuracy to detect much smaller changes and much more accurately than in our previous work^36^.

**Fig. 3.**
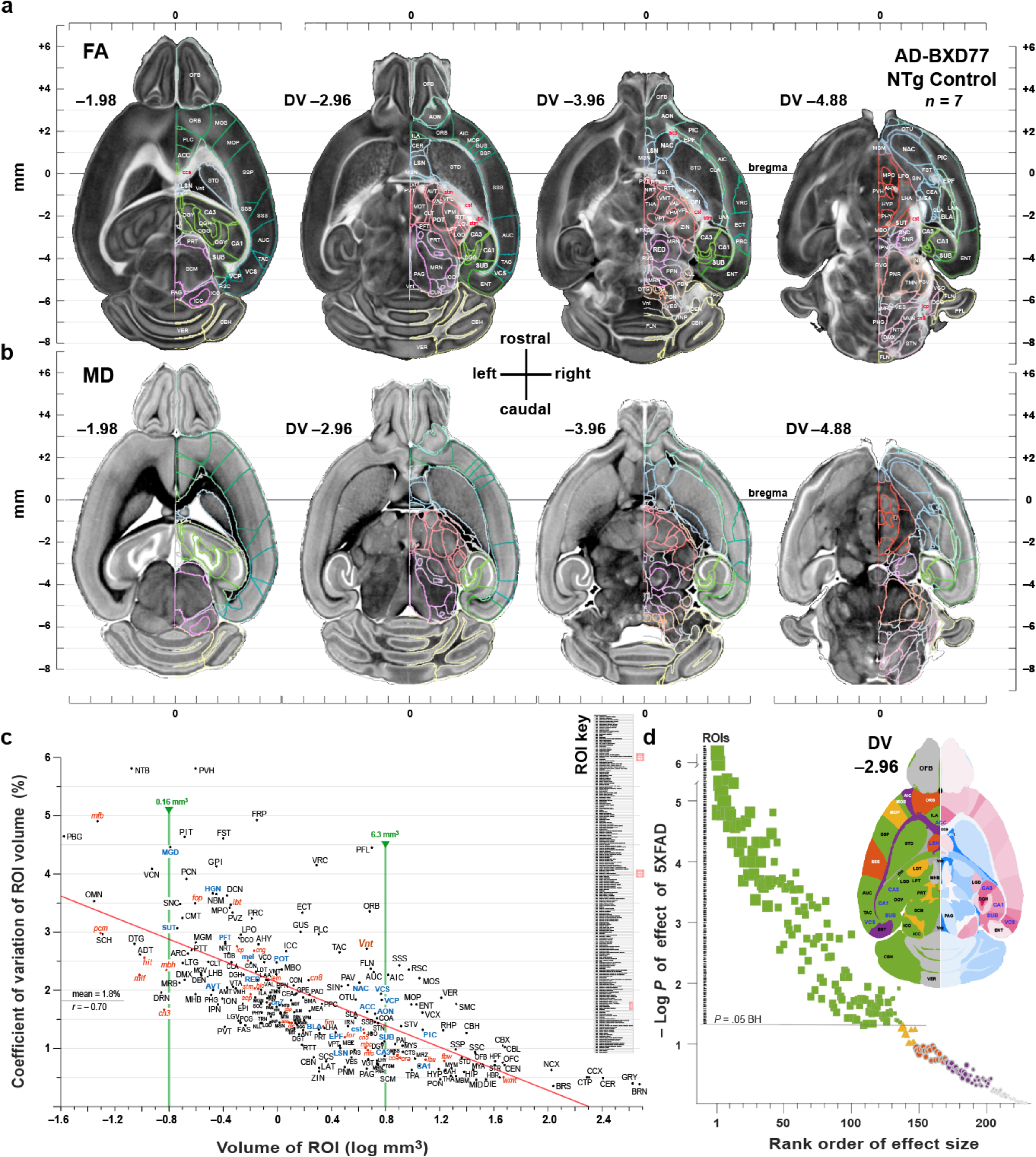
Diffusion tensor delineations of cases and controls are accurate and detect many effects of 5XFAD. MRH data types for a set of AD-BXD77-F1 controls. **a,** Fractional anisotropy at four axial levels of brains of a set of seven AD-BXDF1 controls that have been merged into a common framework using the DMBA. **b.** Mean diffusivity (MD) provides images with good delineation of nuclei. **c,** Volume of regions (bilateral sums) have low average right-versus-left coefficients of variation (1.8%) of ROI volumes across all 57 cases and controls that range from a high of 6% for small structure such as the NTB to a low of 0.5% for whole brain (mean and correlation given to the left near the y axis). The two vertical green lines highlight small (0.16 mm^3^) and larger ROIs (6.3 mm^3^). Those marked in bold blue font along near the green lines were studied using LSM and stereology to estimate amyloid deposition and neuron numbers (Fig. 5) of AD-BXD77 cases and their controls. All structures are labeled using DMBA acronyms. Most nuclei are labeled using black font, and other fiber tracts and the ventricular volume using red font. **d,** Significance of effects of 5XFAD on the volumes of 230 regions. The Y axis gives the –log*P* of the multiple-test corrected *P* value of an ANOVA model of 5XFAD effects using three cofactors: sex, genetic background, and scanner system. X axis is the rank of the Cohen’s D effect size.

There is a strong rostral-to-caudal gradient in the impact of the 5XFAD transgene on gray matter volume. The great majority of ROIs in rostral telencephalic grey matter structures increase in volume—only the caudate-putamen and pallidum are exceptions (**Fig. 3a**). These apparently swelling ROIs include almost all parts of the olfactory system, the entire neocortex, the amygdala complex, and the hippocampal formation. In contrast, almost all diencephalic, mesencephalic and hindbrain structures decrease in volume, as well as most fiber tracts, including the fimbria and fornix (**Fig. 4a,b**). Two notable exceptions from the trend toward shrinkage are the dorsal lateral geniculate nucleus (LGD) and the inferior colliculus (ICC and ICO)—both crucial in sensory processing.

**Fig. 4.**
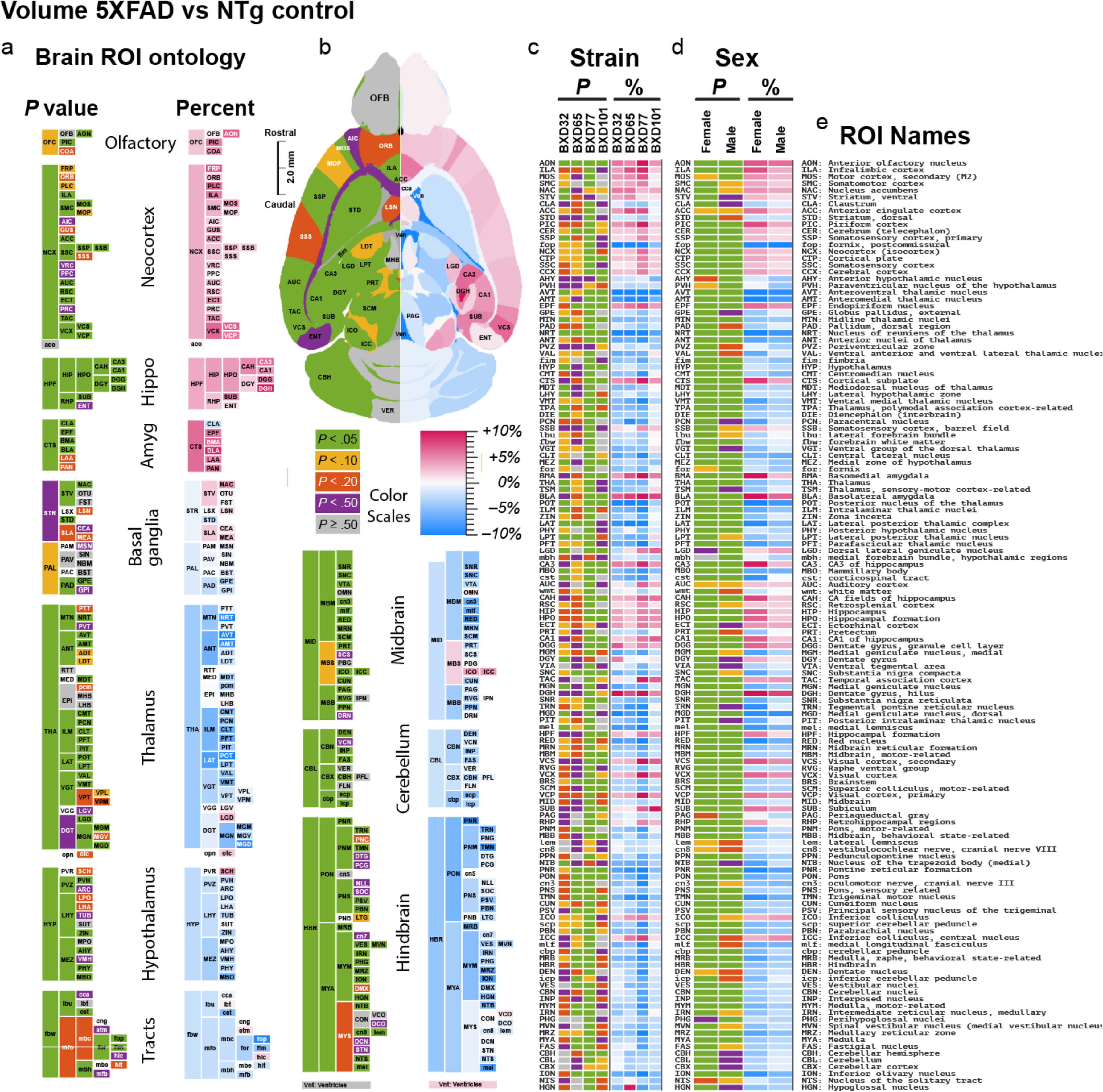
Regional changes in volume associated with Alzheimer’s disease mutations. **a,** Columns of brain regions organized roughly from rostral to caudal as an ontology of stacked colored blocks, each representing an ROI. The colored blocks to the far left define *P* value levels of differences in volume between 5XFAD and control siblings (see Methods). The color scale is defined in the center of the figure beneath the brain section. Green indicates *P* < .05, FDR corrected. Corresponding pink and blue columns to the right in **a** define percentage gains (pinks) and losses (blues) in volume. **b,** Horizontal schematic section of a mouse brain taken from the rCCF3 segmentation of structures (see the *Duke Mouse Brain Atlas of C57BL/6J*). This section is at a level –2.37 mm ventral to the lambda-bregma plane. Color codes as in **a**. The olfactory bulb (OFB) is at the top (rostral) and the vermis (VER) of the cerebellum is at the bottom (caudal). A small number of ROIs are marked for orientation. **c,** Strain genetic background effects for the set of 137 ROIs that achieve a global *P* < .05, now with the significance level (left) and effect size (right) for each of the four AD-BXDF1 types. The genetic background of the sire of each F1 is indicated at the top— BXD32 etc.). **d,** Sex differences and effect sizes for females and males with encoding as in **c**. **e,** A decoding of the three letter acronyms with short descriptions of ROIs that we use in the Duke Mouse Brain Atlas that are derived from CCFv3. The complete list of 231ROIs, their precise *P* values and effect sizes, GeneNetwork.org accession numbers (including right and left sides), and matched Wikipedia entries are provided in *Supplementary Table 1*.

These striking regional changes in volume are corrected for minimal within group variation in brain size and a total of 137 ROIs have significant fraction volumetric changes at *P* < .05 with a Benjamini and Hochberg corrections (**Fig. 3d**, **Fig. 4**). The sizes of the 5XFAD effects are modest in absolute terms, amounting to differences generally less than 10%. While 5XFAD also causes a general decrease in fiber tract volumes, only three tracts reach significance—the corticospinal tract (cst), the fimbria (fim), and post-commissural fornix (fop). The anterior commissure (aco) and the optic chiasm and tract (oct) are the least affected. The total ventricular volume (Vnt) is reduced in volume, this effect is not even close to being significant.

#### The 5XFAD transgenes increase variability in volumes between right and left sides of the brain

Asymmetry between right and left ROI volumes is greater in 5XFAD cases than controls—1.83% ± 0.029 SEM versus 1.72% ± 0.038. This small 6.4% increase could be caused by ontogenetic differences, but we think that a more likely cause is asymmetry in β amyloid deposition that is notable, for example, in figures 5a–d.

**Fig. 5.**
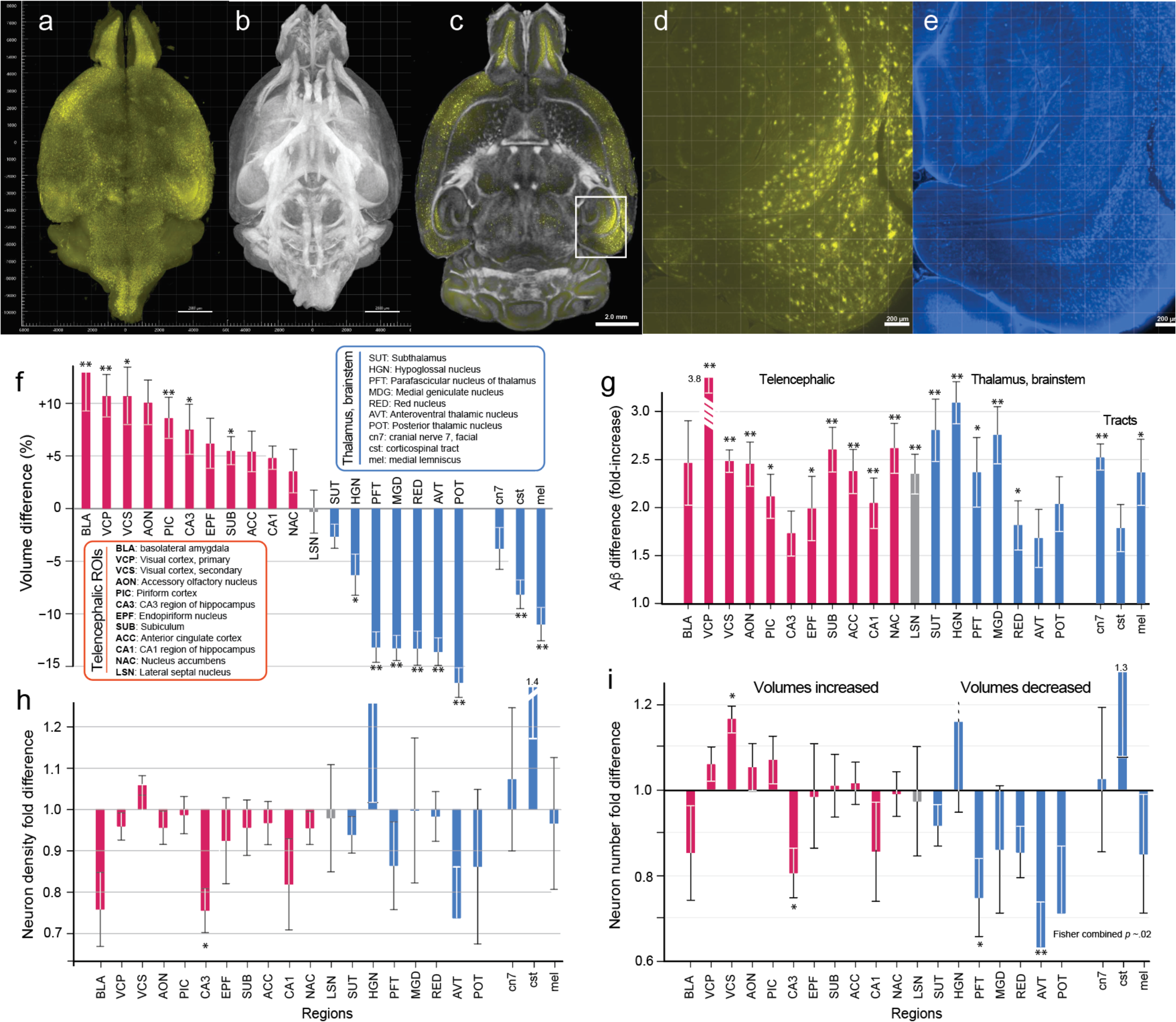
Impact of 5XFAD on aβ aggregates and density of NeuN-positive neuron number. **a,** 3D light sheet microscopy volume-rendered image of β amyloid deposition. **b,** Matched FA volume-rendered image of the same 5XFAD AD-BXD77 F1 brain. **c,** Superposition of β amyloid and FA images demonstrates the accuracy of volume registrations. **d,** Zoomed-in image of the hippocampus (boxed area in **c**) with intense β amyloid in subiculum and entorhinal cortex in the right hemisphere. **e,** NeuN of the same region. **f,** Volume differences of 22 ROIs between AD-BXD77 F1 cases and their controls. ROIs have been ranked from left to right based on the percentage increase or decrease in volumes (left y-axis) with three fibers tracts to the right side. Single asterisk for nominal *P* values <.05, and two asterisks for <.01 (all two-tailed). Volumes were compensated for differences in brain weight among individuals by computing fractional volumes of ROIs given whole brain volume. The basolateral amygdala (BLA) to the left side has the greatest percentage increase in volume in the 5XFAD cases; the posterior nucleus of the thalamus (POT) has the greatest percentage decrease in volume in the 5XFAD cases. **g,** Amyloid β aggregates are 1.7-to 3.8-fold higher in 5XFAD cases than controls. **h,** Neuron densities are lower in 10 of 11 regions that have increased volumes in 5XFAD cases, but only the CA3 has a nominally significant reduction in density. Neuron density changes in ROIs that are smaller in 5XFAD cases (blue) are generally noisy due to their much smaller volumes (Fig. 3c). **i,** Neuron numbers are reduced in CA3, PFT, and AVT at nominal *P* values of <.05. The apparent increase in neuron numbers in secondary visual cortex (VCS) of 5XFAD cases is likely to be an artifact caused by interactions between protocols and β amyloid deposition and cellular stress (see text). NeuN counts in fiber tracts were very low on an absolute scale, and the fold changes are likely to reflect low-level mislabeling and erroneous counts. We computed the Fisher’s combined *P* value for the 7 ROIs with volumetric reduction (tracts excluded) and numbers are collectively lower in 5XFAD cases (Fisher’s combined *P* of .02). All stereology from left sides only.

#### Strong sex effects of the 5XFAD transgenes

We initiated this analysis by testing our ability to resolve the well-known sex difference in the volume of the medial preoptic area. The bilateral volume of the MPO is 0.428 ± 0.033 SD mm^3^ in females and 0.468 ± 0.024 mm^3^ in males—about a 9% difference. In an ANOVA fitting strain, transgene status, sex, the sex-by-transgene interaction, only sex is a strong predictor of variance (F = 38.3, *P* < .0001). Of the 137 ROI highlighted in the overall n-way ANOVA as being modulated by transgene status, at least 28 have significant sex differences at a Bonferroni criterion of *P* < .05 (**Fig. 4d** and *Supplementary Table 6, column 7*). In contrast, and somewhat surprisingly, not a single ROI has a significant 5XFAD-by-sex interaction).

#### Genetic background strongly modulates severity of 5XFAD effect

Genetic background is a highly significant predictor of variance in 5XFAD transgene action on the fractional volumes of ROIs that correct for overall differences in brain size (*F* = 6.60, *P* < .0008). Thirty-one ROIs have significant differences in fractional volume even with aggressive Bonferroni correction at *P* < .05 (**Fig 4c**, *Supplementary Table 5, column 6*). By far the strongest effect of the 5XFAD transgene in on the AD-BXD77 F1 background. However, the trends of effects of all four strains tend in the same direction and there is no significant 5XFAD-by-strain interaction.

### 2. Light sheet stereological analysis of changes in NeuN+ cell densities and numbers in AD-BXD77 cases and controls (see *Supplementary Table 2* for acronyms)

The volumes of 11 of 12 telencephalic ROIs are larger in AD-BXD77 F1 cases than in their sibling controls (left side of **Fig. 5f**). This difference is nominally significant for half of all regions. The Fisher’s combined *P* value is highly significant (*P* < .0001). This pattern is replicated, but with the opposite polarity, for all thalamic and brainstem structures. Figure 5g and h address the question of whether there is evidence in AD-BXD77 cases of loss of NeuN-positive cells. Gurdon and colleagues estimated the intensity of NeuN staining in 6- and 14-month-old AD-BXD F1s and detected notable loss of intensity in the hippocampus proper (CA1–CA3), in the dentate gyrus, in the posterior amygdala nucleus (PAN), and in the striatum-like amygdalar nuclei (the central nucleus of the amygdala and parts of the medial nucleus of the amygdala)^10^. We have counted neurons in both CA1 and CA3, and in both we detect appreciable loss of NeuN+ cell densities—about 20% in both cases that is neatly matched to absolute loss of numbers (Fig. 5h). In the case of CA3, the loss in *density of cells* is nominally significant (*P* =.013, two-tailed), but the loss of *numbers of cells* does not reach this level (*P* = .076, two-tailed). Given that the null hypothesis is polarized (no loss of neurons), a one-tailed test is justifiable. Our best estimate is that by 14 months about 20% of NeuN+ cells have died in CA3 of AD-BXD77 F1s.

We also have evidence of cells loss, especially in smaller thalamic, mesencephalic, and hindbrain structures that have not been well studied in either mouse models of AD or in human AD patients. In six of seven ROIs that have smaller volumes in 5XFAD cases there is also a significant reduction in cell numbers using Fisher’s combined *P* test (*P* of .017 two-tailed). Numbers are 35% lower in the anteroventral thalamic nucleus (AVT, *P* = .009, two-tailed), and 25% lower in the parafascicular thalamic nucleus (PFT, *P* = .044, two-tailed).

There is one finding we struggle to explain—the apparent increase in NeuN+ cells in the secondary visual cortex (**Fig. 5i**, VCS). Cases have an average population of 363,000 ± 8200 SE cells versus 311,000 ± 9500 SE in controls (*P* = .006). We have reviewed the delineations of VCS and recounted all cases and controls, and we did not identify any operational errors. Busser and colleagues^37^ discovered that AD in humans can induce aberrant expression of cell cycle genes due to cellular stress, and the RBFOX3 protein to which the NeuN antibodies normally bind is known to be involved in both cellular stress, as well as linked to cell cycle proteins in the nucleus^38^. It is therefore possible that ectopic or artifactual NeuN+ labeling leads to increased detection of NeuN+ cells. Finally, NeuN can cross-react with synapsin1 protein^39^ and is also expressed in GFAP-positive astrocytes in cell culture^40^.

### 3. Behavioral differences covary with MRH ROI volumes

We have overlapping behavioral and neuroimaging data for 52 AD-BXD individuals—31 cases and 21 controls; 27 females and 25 males. We focused attention on contextual fear acquisition (CFA) and contextual fear memory (CFM). We computed correlations of these two traits with variation of all 230 ROI volumes (bilateral sums, whole brain excluded).

Correlations of both the hilus of the dentate gyrus (DGH) and the posterior amygdala (PAN) with contextual fear memory are negative (**Fig. 6b and 6d**). In other words, paradoxically, the larger these ROIs, the worse an individual’s memory retention. In contrast, figure 6a highlights a positive correlation of *r* = 0.49 between performance and greater volume in the fimbria (fim) with acquisition, and in figure 6f a positive correlation of *r* of 0.49 of the dorsal thalamus (THA) volume with memory. All correlations are significant (*P* < .05) whether using Pearson or Spearman methods and are also corrected for multiple tests (*Supplementary Table 8*).

**Fig. 6.**
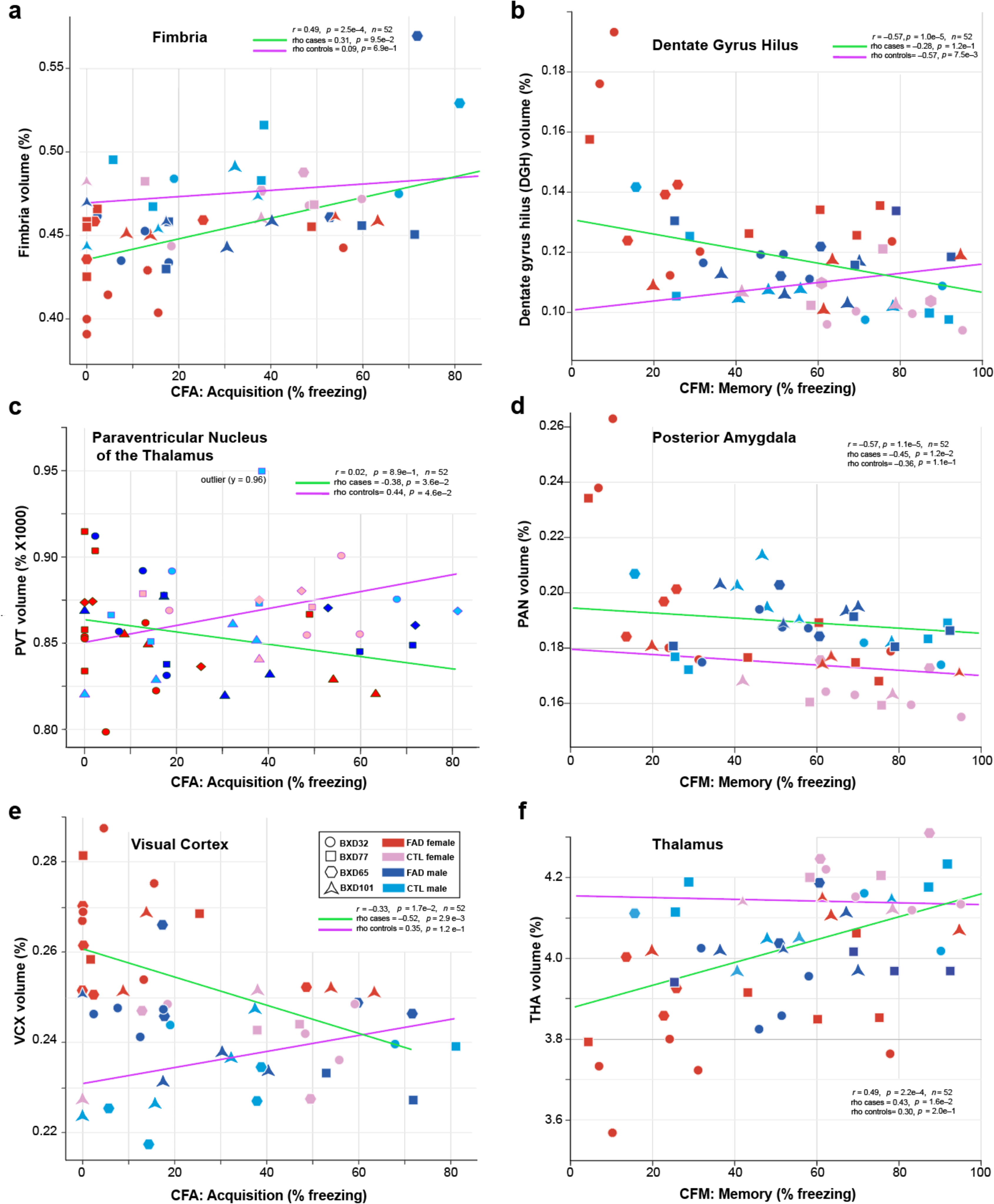
Volume changes in 5XFAD cases are linked to worse behavioral performance. **a,** Volume of the fimbria is smaller in cases than controls and fear acquisition is generally worse. Pearson correlation lines for cases (green) and for controls (purple) are given with the *combined r* correlation with P values in each panel. We also provide the rank order rho correlations separately for case and controls. **b,** Volume of the hilus of the dentate gyrus is larger in 5XFAD cases than controls. The 5XFAD cases have higher mean Y axis values and worse contextual fear memory (lower percentage freezing). **c,** Volume of the paraventricular nucleus of the thalamus correlates with slightly worse CFA in cases than controls. This ROI is so small that the y axis scale has been modified and a true value of 0.00028% is labeled 0.28. One outlier measurement is flagged. **d,** Volume of the posterior amygdala in 5XFAD cases is greater but there is minimal impact on CFM. **e,** Volume of the visual cortex is negatively correlated with CFA in cases but is positively correlated in controls. **e,** Volume of the VCX is negatively correlated with CFA in cases but not controls. **f,** Volume of the thalamus is decreased in 5XFAD cases, but there is no significant difference between cases and control either by Pearson correction (lines) or the rho correlations (text values in panels).

A more interesting question is whether there are differences in correlations between the two groups—5XFAD cases and their controls. We therefore stratified the analysis by transgene status (*Supplementary Table 8*) and tested for significant differences between rank-order correlations for all ROIs with both contextual fear traits— CFA and CFM. A test of this sort with modest sample sizes (*ns* from 21–31) is not well powered. Nonetheless, we detected 46 ROIs for which correlations between cases and controls have opposite polarities that are nominally significant with fear acquisition—a measure of learning efficacy (e.g., **Fig. 6c,e**). None meets FDR corrections at *P* < .05. Remarkably, the visual cortex (Fig. 6e) has among the highest differences in correlations between cases and controls for both CFA and CFM. Both temporal association cortex and the paraventricular nucleus of the thalamus also differ markedly (**Fig. 6c**, PVT where rho is –0.38 for cases and +0.44 for controls).

In contrast, we detected only three structures with differences for CFM in correlations between cases and controls—the medial septal nucleus (MSN, rho of +0.40 for cases vs –0.29 for controls), the visual cortex (VCX, rho –0.41 cases vs +0.18 for controls), and the ventromedial hypothalamic nucleus (VMH, rho +0.11 for cases vs –0.45 for controls). The detection of a large difference in correlations in the MSN is particularly interesting given its critical role in CFM via modulation of CA1 function^41,42^. The detection of differential correlations with visual cortex is also intriguing given its critical role in contextual fear memory via modulation by microglial tone ^43^.

Given the large number of tests, even the strongest results will benefit from larger sample sizes. Nominal and FDR-corrected *P* values are provided for all 230 ROIs for both CFM and CFA in *Supplementary Table 8*. Rather than making claims about these correlations we prefer to regard them as proof-of-concept that support the hypothesis that we are technically able to detect correlations between ROI volumes, changes in neuron densities and numbers, and changes in specific behaviors known to be diagnostic of features of human AD. The apparent link to visual system ROIs is surprising but is consistent with a prodromal form of posterior cerebral atrophy (PCA) type of early onset AD in humans^44^.

## DISCUSSION

Until recently the most definitive diagnosis of human AD has involved specific patterns of cognitive decline, often corroborated by postmortem evaluation of amyloid aggregates and loss of brain mass and neuron numbers^45,46^. Over the last 25 years MRI has become one of the more widely used methods for diagnoses of AD subtypes^47,48,44,45,49^. Initial MRI diagnoses of AD has focused on morphometric measures of global and regional CNS atrophy. Spin lattice (T1) and spin-spin relaxation (T2) have been exploited to estimate loss of myelin integrity. Diffusion tensor imaging (DTI) has added additional quantitative indices of tissue cytoarchitecture—the DTI scalar diffusion metrics, namely axial diffusivity (AD), mean diffusivity (MD), radial diffusivity (RD), and fractional anisotropy (FA). DTI can be extended to a systems level by the computation of whole brain connectomes that provide global evaluation of structural connectivity. In combination with advanced statistical methods, it is now practical to link volumetric changes and those of the structural connectomes to behavioral decline.

Even the earliest-onset familial forms of Alzheimer’s disease with well-known genetic causes^50,51,52^ are complex in terms of their progression and symptomatology. For example, highly penetrant mutations of *PSEN1* are associated with a 30-year range of onset even in single families^53, 54,55, 56^. There are also striking differences in the range of behavioral deficits with brain regions that either swell or atrophy^57,58,59^, including posterior cortical atrophy (PCA)—a relatively common variant of AD (∼10% of early onset cases) that causes a wide range of visual and other function deficits^60,44,61,62,63^. This complexity has many causes, but two are critical: genetic ancestry and sex. In the present work we have addressed both genetic background and sex and have detected pervasive effects of human AD mutations on volumes of 137 of 230 well defined cortical and subcortical brain regions in mice, including in nuclei as small as the perifascicular nucleus (0.21 mm^3^ unilateral, **Fig. 3c,d**, **Fig. 4**, **Fig. 5**).

One major finding of our work is a strong rostral-to-caudal gradient in effects of the 5XFAD transgene that causes an increase in volumes in most of the telencephalon and cortex, but a shrinkage of almost all other regions of the brain. We also corroborate that there is a loss of neurons in the CA3 of the hippocampus in the most severely affected AD-BXD77 cases. While not yet sufficiently robust to make strong claims, we suspect that there is appreciable neuron loss also in anterior and midline nuclei of the thalamus on this genetic background. Our results also confirm and extend previous work by Kaczorowski and colleagues that the effects of 5XFAD are highly variable as a function of genetic background^10,12^. We confirm generally stronger effects of AD in females that is also a consistent characteristic of FAD in humans^64,65^.

Our strong positive findings are in contrast to negative findings of the MODEL-AD consortium^24^ and several other recent studies^9,66^ using both conventional T1/T2 MRI and DTI (overview of these papers in Tables 2 and 3). One possible biological explanation for some failures is that the impact of 5XFAD has often been studied only one the C57BL/6J genetic background—a background known to be resistant to the effects of this transgene^12^. But three key technical explanations are even more important: 1. the higher resolution of the *ex vivo* MR images we have generated (**Fig. 2a,b**); 2. the use of more sophisticated multiangle DTI methods that enhances contrast and the precision of delineations; and, 3. the much larger number ROIs that we have been able to delineate.

**Table 1.** Numbers of cases and control classes.

| Strain<br>of Sire* | 5XFAD Tg<br>F1 (n) # |  | NTg CTL<br>F1 (n) # |  | Total |
| --- | --- | --- | --- | --- | --- |
|  | M | F | M | F |  |
| BXD32 | 4 | 5 | 2 | 5 | 15 |
| BXD65 | 2 | 4 | 1 | 2 | 9 |
| BXD77 | 4 | 6 | 5 | 2 | 17 |
| BXD101 | 5 | 4 | 4 | 3 | 16 |
| **Total** | **14** | **20** | **12** | **11** | **57** |

\* Type of BXD strain crossed to C57BL/6J 5XFAD congenic carrier

**Table 2.** Recent MRI studies of rodent models of AD using T1/T2 weighted MRI.

| Reference | Transgene | N of Animals | In- or Ex-Vivo | Field (Tesla) | Voxel (mm<sup>3</sup>) | N of ROIs |
| --- | --- | --- | --- | --- | --- | --- |
| Badea, Johnson et al. 2010 (Ref <sup>67</sup>) | APP/TTA<br>TTA | 6<br>6 | Ex | 9.4 | 0.000011 | 33 |
| Spencer, Bridges et al. 2013 (Ref <sup>68</sup>) | 5XFAD | 6 Tg<br>6 NTg | In | 4.7 | 0.054000 | 3 |
| Girard, Jacquet et al. 2014 (Ref <sup>69</sup>) | 5XFAD | 30 Tg | In | NA | 0.003300 | 3 |
| Micotti, Paladini et al. 2015<sup>70</sup> | TASTPM<br>APP/ Tau | 30 Tg<br>40 Tg | In | 7T | 0.002500 | 2 |
| Spencer, Lovell et al. 2017m (Ref <sup>71</sup>) | 5XFAD | 11 Tg<br>10 NTg | In | 4.7T | 0.054000 | 3 |
| Frost, Longo et al. 2020 (Ref <sup>72</sup>) | 5XFAD | 6 Tg<br>6 NTg | In | 7T | 0.014000 | 6 |
| Oblak, Lin et al., 2021 (Ref <sup>24</sup>) | 5XFAD | 58 Tg | In | 3T | 0.006500 | 56 |
| Zhou, Ngiam et al. 2022 (Ref <sup>73</sup>) | 3XTg | 10 3xTg<br>7 NTg | In | 9.4T | 0.003000 | 1 |
| Kim et al. 2021; Kim et al. 2023 (Refs <sup>74,75</sup>) | 5XFAD<br>TgF344- rat | 8 Tg<br>6 NTg | In | 9.4T | 0.000900 | 2 |
| Tomaszewski, Sukstanskii et al. 2024 (Ref <sup>76</sup>) | Tg4510 | 10 | In | 7T | 0.004200 | 1 |

**Table 3.** Representative MRI/DTI studies of mouse models of AD.

| Reference | Transgene | N Animals | In- or Ex-vivo | Field (T)<br>b (s/mm<sup>2</sup>) | Voxel (mm<sup>3</sup>)<br>Relative Resolution * | N of angles | N of ROIs |
| --- | --- | --- | --- | --- | --- | --- | --- |
| Grandjean, Derungs et al. 2016 (Ref <sup>77</sup>) | ArcAβ, E22ΔAβ, PSAPP | 33 Tg, 35 NTg | In | 9.4<br>1000 | 0.0059<br>320 | 36 | 6 |
| Kesler, Acton et al. 2018 (Ref <sup>78</sup>) | 5XFAD | 4 Tg<br>4 NTg | Ex | 9.4<br>1000 | 0.00083<br>53 | 20 | 32 |
| Badea, Wu et al. 2019 (Ref <sup>79</sup>) | APOE4, APOE3 | 10<br>14 | Ex | 9.4<br>4000 | 0.00017<br>11 | 46 | 332 |
| Falangola, Nie et al. 2021 (Ref <sup>80</sup>) | 3XTg-AD | 89 Tg<br>56 NTg | In | 7<br>2000 | 0.017<br>1088 | 30 | 3 |
| Falangola, Nie et al. 2022 (Ref <sup>80</sup>) | 3XTg-AD | 13 Tg, 11 NTg | In | 7<br>2000 | 0.017<br>1088 | 30 | 2 |
| Murdy, Dunn et al. 2022 (Ref <sup>9</sup>) | 5XFAD | 96 BXD | In | 7<br>1200 | 0.020<br>1280 | 46 | 83 |
| Maharjan, Tsai et al. 2022 (Ref <sup>81</sup>) | 5XFAD | 8 5xFAD<br>6 NTg | Ex | 9.4<br>3000 | 0.0010<br>64 | 61 | 166 |
| Westi, Molhemi et al. 2024 (Ref <sup>66</sup>) | 5XFAD | 7 5xFAD<br>7 NTg | Ex | 9.4<br>2000 | 0.0045<br>288 | 120 | 8 |
| Winter, Mahzarnia et al. 2024 (Ref <sup>82</sup>) | APOE2, APOE3, APOE4 | 173 | Ex | 9.4<br>2000<br>4000 | .000091<br>6 | 51 | 332 |
| This Work* | 5XFAD | 57 BXD | Ex | 9.4<br>3000 | .000016<br>1 | 61 | 308 |

\* Resolution is normalized to the present study. A relative resolution of 320 means voxels in that reference are 320 times larger. *In vivo* studies are 320 to 1984 times lower resolution than this work. *Ex vivo* studies with relative resolution DWI noted in Figures 2 and 3 are made possible by the large number of angles ( $n$ of 61) we have sampled.

The increase in volumes of ROIs in cortical regions has, to the best of our knowledge, never been reported in humans with either *PSEN1* or the *APP* mutations. Quite the opposite—a common denominator of FAD is cortical thinning even in presymptomatic individuals^83,84^. We consider some technical factors below but given the consistency of the apparent swelling of most telencephalic regions and the ease with which many of these large ROIs can be delineated, we currently hypothesize that the increased volume is an initial and perhaps an inflammatory response—potentially compensatory—of these ROIs to beta amyloid aggregation—a finding supported by the work of Andersen, Westi, and colleagues^85,66^. Whatever processes are involved in olfactory, cortical, and hippocampal ROI volume increases, they are not shared by thalamic, hypothalamic, and brainstem regions. The regional specificity is a major clue and constraint on potential models of this rostral-to-caudal gradient of 5XFAD effects in AD-BXD F1 mice.

### Sex differences in the impact of 5XFAD on regional brain volume

The core of what we need to say on this important topic is summarized in figure 3d. Of 137 ROIs with volumes that are significantly affected by the 5XFAD transgene, only nine have stronger effects in males than in females (MOS, AHY, PVH, for, LGD, mbh, PAG, PHG, and NTS), and many of these apparent differences may be due to sampling error. However, this list does highlight hypothalamic and other midline regions for scrutiny as sex-dependent targets of the human *APP* and *PSEN1* mutations^8, 64, 83^. (See *Supplementary table 2* for all ROI acronyms.)

### Relationships of β-amyloid burden and NeuN+ cell densities and numbers

We demonstrate significant volumetric changes across brain regions in AD-BXD F1 cases—especially on the AD-BXD77 hybrid background. This striking regional variation in volumetric effects leads to several questions about underlying molecular and cellular processes. We can provide some initial insights into the quantitative cellular correlates, but we cannot yet provide clear answers.

The autosomal dominant mutations in *APP* and *PSEN1* are known to cause aberrant APP cleavage and clearance^86,6^ and this leads to neuroinflammation mediated by the activation of astrocytes and microglia, as well as fundamental changes in cortical and hippocampal bioenergetics^85,66^. We detect what could be considered parenchymal swelling in many telencephalic ROIs. This increase in volume in cortex initially puzzled us, but there are strong precedents in human neuroimaging studies, particularly in presymptomatic *PSEN1* humans in which several cortical areas are initially approximately 250 µm thicker than control but are 250 µm thinner in when symptomatic (see figure 5 of Ref^57^). ROI volumes also initially increase in asymptomatic cases relative to controls, but this is followed by severe loss in symptomatic cases. These *PSEN1* subjects carried either M139T, K239N, or L286P mutations. The 5XFAD transgene incorporates the L286V mutation, and to add insult to injury, also incorporates the M146L mutation.

The increase in volume of the hippocampus is more than “prodromal” and can be linked to significant loss of neurons. This is most evident in CA3, and perhaps also in CA1 (**Fig. 4i**). Here our results align with the work of Gurdon and colleagues^10^ who detected a decrease in NeuN+ intensity in the hippocampus. We estimate that 10–20% of pyramidal neurons are lost in AD-BXD77 F1 cases. To the best of our knowledge this is the first, stereological estimate of absolute neuron loss in 5XFAD mice (not just a reduction in density). In sum, this work suggests that the increase in volume of some ROIs is a prodromal stage of disease that can be associated, or not, with neuron loss. A surprising coda is that an increase in the volume of the hippocampus (+7% in CA3 of AD-BXD77 F1 cases) is associated with a significant loss of neurons (∼18%).

The light sheet analysis can also dissociate amyloid levels from the polarity and amplitude of volumetric change (**Fig. 5f,g**). Amyloid burden is high across all 22 ROIs in AD-BXD77 cases, but this high expression does not covary with volume change (rho = +0.27, *P* = 0.2). This is also true of many other ROIs. Gurdon and colleagues^10^ demonstrate that both astrocytes and microglia are activated in AD-BXD F1 mice in *all* brain regions, including striatum, pallidum, thalamus, and brainstem (Ref^10^, their figure 4). In our work, these same regions have reduced ROI volumes—a clear dissociation between volumetric changes and classic markers of neuroinflammation.

If molecular drivers of AD ultimately converge on neuroinflammation, then we need better causal explanations for spatial heterogeneity. Many differences in expression, cell types, and patterns of activity might explain volume increases and decreases and neuron loss or sparing. We also need to explain both the broad rostral-caudal gradient in swelling and shrinkage, as well as the sharp differences between neighboring ROIs. One concrete example may help make this point—the dorsal lateral geniculate nucleus (LGD) is the only thalamic volume that is larger because of the 5XFAD transgene at 14 months of age. However, its immediate neighbor, the lateral posterior nucleus gets smaller (LPT, this is the homolog of the human pulvinar, see Fig. 4b). Both nuclei are crucial components of the visual system, and both have dense reciprocal connections to the visual cortex—both VCP and VCS. But these two thalamic nuclei have radically different physiology and function. It is intriguing that of all nuclei in the dorsal thalamus of mice, the LGD has by far the highest concentration of GABAergic interneurons^87^.

Two thalamic ROIs with significant volumetric losses also lose many neurons in AD-BXD77 mice—the perifascicular nucleus (PFT) and the anterior ventral nucleus (AVT) (Fig. 4i). Both nuclei have roles in cognition and memory. The anterior nuclear group, including the AVT, have well known roles in memory ^88^. The thalamus is one of the first regions to have high amyloid levels and volumetric atrophy in presymptomatic carriers of *PSEN1* mutations^89^. Given our strong signal in anterior and midline thalamic regions, these regions are worth more intense study as early targets of AD.

### Linkage of ROI volumes with behavioral variation

Even in the relatively small subset of animals in which we have both neuroanatomical and behavioral data, we see significant correlations with FDR correction. A remarkable set of 29 regions are correlated with CFA—a key learning metric. Twelve of these ROIs are thalamic (AVT, ILM, PCN, PFT, PIT, POT, THA, TPA, TSM, VAL, VGT, VMT), 2 are hypothalamic (SUT, ZIN, lbu), 2 are tracts associated with hippocampus and many other ROIs (for, fim), 1 is the basomedial amygdala, and the remaining 7 are motor-related (ION, MRN, MBM, PNM, PRT, RED, TMN, cst). Collectively this set highlights many linked sensory, perceptual, fear and affect, and motor integration that will be critical in fear conditioning^90,91,92,93^

With respect to CFM, the data highlight both the entire dorsal thalamus (THA) and 3 of its components (LAT, TPA, DGT), 3 amygdalar regions (PAN, LAA, BLA), the hilus of dentate gyrus (DGH) and the cortical subplate (CTS)^94^. All of these ROIs have been previously associated with fear memory^95,96,97,98,99,92,100,101,102^ and this gives us confidence in the validity of both methods and our findings.

Volumes of the whole thalamus (THA) and the polymodal association cortex-related part of the thalamus (TPA) correlate positively with both CFA and CFM, indicating linkage with both learning (acquisition) and memory. This is in good agreement with much work done at the circuit level^98^. That we see the thalamic regions so prominently is important, because these regions are being increasingly recognized as important in AD^88,103^and reduced volume of several thalamic regions have been associated with both early- and late-onset disease^104^.

Several hippocampal and amygdalar ROIs have positive correlations – that is, larger volumes are associated with worse function—against naive expectations^105,106,107^. One explanation mentioned above is parenchymal swelling early in AD progression, and a possible compensatory mechanism^108,109^.

However, the above correlations represent processes taking place collectively across both cases and controls. Of greater interest are the subset of correlations that are significantly different between cases and controls. We identify 56 ROIs for which correlations differ in directions or magnitude for CFA, and 13 for CFM. Of these, 11 are nominally significant for both CFA and CFM (VCX, VCP, PSV, VCS, ECT, pcm, MGD, RSC, mlf, PTT, cn3). Five of these are involved considered sensory or perceptual (VCX, VCP, PSV, VCS, MGD), and deficits of these types are often symptom of AD, often occurring before the more diagnostic cognitive complaints^110 111,112,113^. Three other ROIs (ECT, RSC, PTT) have been previously linked to learning and memory function. Finally, two tracts—the medial longitudinal fascicle (mlf) and the central part of the oculomotor nerve (cn3) are both involved with eye movement control. Oculomotor cranial neuropathy has recently been identified as an early indicator of AD^114^.

### Preclinical studies of mice require high resolution to detect 5XFAD effect on brain structure

Preclinical protocols that parallel clinical MRI have obvious benefits because they enable much tighter control of genetics, age, diet, and treatments. Unfortunately, most preclinical studies of AD have been undertaken using a single strain of mouse, and this severely limits translational relevance of results to clinical populations. The second problem is that brains of mice are roughly 3000 times smaller than those of humans. Scaling spatial resolution to achieve the same voxel density has been challenging. But increasing spatial resolution is not the only requirement. It is also essential to be able to accurately measure physical changes caused by mutations and transgene on brain structure at the level of discrete ROIs. There are multiple candidate measures: volume changes, changes in diffusion properties, changes in connectomes and/or network properties (derived from diffusion images). Previous work in mouse models of AD (Table 2) has generally used simple T1 and T2 weighted images to measure structural differences and changes in relaxation times.

Table 3 summarizes some of the more recent preclinical studies using DTI. These studies compare diffusion properties, more specifically FA and MD, or connectomes and their network architectures. In both tables we provide spatial resolution metrics in absolute voxel volume (mm^3^) and as relative resolution normalized to the 25 µm^3^ resolution that we are now using routinely.

Data for the two previous high-resolution studies^79,82^ were acquired in the CIVM at Duke using our earlier protocol ^115,36^. Both focused on changes in diffusion and network properties but did not measure ROI volume changes.

Kelser and colleagues studied eight individuals at an isotropic resolution of 0.094 mm (voxels of 0.00083 mm^3^) with b = 1000 s/mm^2^ and with 20 angular acquisitions^78^. Their analysis of structural connectomes between 32 ROIs showed no differences between cases and controls.

Westi and colleagues describe a comprehensive analysis of the 5XFAD model employing functional MRI, diffusion MRI, and immunohistochemistry^66^. They employed a multi-shell dMRI sequence to measure diffusion kurtosis, a second-order measure of diffusion properties in eight manually delineated ROIs. They also detected no significant difference in metrics between cases and controls, and no differences in volumes for any ROI at 2 and 6 months—perhaps too early in disease progression.

The study by Murdy and colleagues^9^ is most closely aligned with the present work. This in vivo structural MRI study included a very high level of genetic diversity among cases and controls and included balanced sets of sexes. Their sample size was unusually large—a total of 90 cases and controls. They also used an innovative NODDI protocol—a more nuanced model of DTI and delineated a set of 83 ROIs derived from the Allen Brain Atlas. Their primary results highlighted many differences in NODDI parameters dependent on genetic background, but they only detected one difference caused by 5XFAD on axial diffusivity signal in the stria medullaris (STM) of cases relative to controls. As noted by the authors, the minimal impact of the 5XFAD transgene in this study is probably due to the early age of imaging (90 days). Of particular interest, these authors also probed the relationship between changes in CFA and CFM and the MRI metrics (their Figure 9) and they also noted strong correlations with visual cortex (their area prostriata is equivalent to our VCS).

Hippocampal and cortical atrophy were noted as some of the earliest clinical biomarkers for AD^116,117,118,119^ using T1, T2 weighted MRI. DTI has yielded additional opportunities to track changes in tissue microstructure through diffusion scalar measures of FA and MD and connectomes and network analysis derived from tractography. Replication of these results in preclinical settings has been limited. We suggest that the problems in preclinical reproducibility stem from radical differences in methods. For example, spatial resolution in clinical MRI varies across studies at most by < 10 (Ref ^120^) but the results in tables 2 and 3 for preclinical studies vary over a much wider range. Methods for reproducible processing in clinical MRI have been routine for nearly 20 years^121^. But application of these methods (over this wide range of variation) have not been as thoroughly validated.

### Caveats and limitations of study

Expression of the bigenic 5XFAD transgene (Tg6799) is driven by neuron-specific *Thy1* cassette vectors ^122^, and this raises the possibility that rostral-to-caudal effects we have detected are downstream of regional differences in promotor activity. This can probably be ruled out because many teams have shown strong expression of the 5XFAD transgene and high levels of amyloid deposition in the thalamus and midbrain^122^— regions in which we have demonstrated highly significant reductions in volume in 5XFAD cases (see Ref^23^, their figure 4; and Ref^24^, their figure 5). Furthermore, the volume reduction in diencephalic, midbrain, and hindbrain regions is just as strong and just as ROI-specific as volume expansions in most telencephalic structures. The only major brain region that appears to equivocate in its responses to the transgene is the striatopallidal complex, in which ventral striatum (e.g., NAC) increases in volume but globus pallidus externa (GPE) decreases in volume—in both cases with significant *P* values.

Our analyses of cell densities and numbers in 3D light sheet image data sets are of course subject to all of the usual caveats associated with immunohistochemical protocols when applied to stereology; primarily issues related to penetration and specificity of NeuN antibodies, non-specific background signal, and in our case to possible interactions of these factors with the consequences of the *APP* and *PSEN1* mutations on tissue integrity and on epitope levels and stability. Nonetheless, the numerical analyses are relatively robust in our opinion and in both cases and controls our count align reasonably well with neuron numbers (also see Tian et al., 2023). However, the high CVs of cell counts will compel us to increase samples sizes by at least a factor of two in order to provide stronger evidence for or against neuronal losses within specific ROIs.

For our most ambitious work of linking ROI volumetric changes to specific alterations in behavior, the main limitation is sample size. We rely on correlations within the cases and the controls and then we estimate the statistical significance of the difference between correlations, region by region. This is an inherently hard problem and to achieve phenome-wide significance across 231 ROIs and nearly 100 traits of interest (TOIs) these will require on the order of 100–200 individuals in each group to be able to gain sufficient power to reach down to FDRs of under 5%. However, with our current data set we are close to having sufficient power to detect differences in ROI-to-TOI correlations between cases and controls with an FDR of 10%.

### Future directions

Perhaps most obviously—we need to carry out similar statistical analyses of key scalar MRI metrics (FA, MD, AD, RD, etc.) and of connectome. These metrics have great promise as neuroanatomical biomarkers of the progression and treatment of AD^29,9^ This work is already in progress using the same set of cases and controls.

Given the success and throughput of the new MRH methods, another obvious direction is to do more of the same. Our main motivation is two-fold: to gain statistical power and to more precisely define architectural changes caused by FAD mutations as a function of age, sex, background, and treatments; and second, to establish causal links between changes at brain systems to degraded behaviors of many types. Both goals are shared by both clinical and preclinical neuroscience research communities.

We add one more atypical goal—namely to increase numbers of AD-BXD F1 types so that we can use quantitative genetic methods to define causal loci that account for susceptibility and resistance to neurodegenerative diseases^123,124,21,125,126^. We currently have MRH data for additional 400 mice with matched behavioral data, but for well powered mapping studies we will probably need data for 40 AD-BXD F1 types and 640 individuals—a heavy but necessary lift.

Finally, it is already practical to use AI-driven methods to refine and extend ROI delineations to ever smaller and more problematic CNS structures. For example, we do not yet delineate CA2 as a discrete ROI, but as suggested by figure 2g, this should soon be possible. We should anticipate a doubling of the numbers of well delineated ROIs.

## METHODS

### AD-BXD F1: a genetically diverse replicable isogenic model

C57BL/6J congenic mice (B6.Cg-Tg(APPSwFlLon,PSEN1*M146L*L286V)6799Vas/Mmjax; RRID:MMRRC_034848-JAX), were obtained from the Mutant Mouse Resource and Research Center (MMRRC) at The Jackson Laboratory^122^. These mice are hemizygous for the dominant 5XFAD transgenes, ^127^ which consists of five human missense mutations that cause familial AD: three in the amyloid precursor protein gene (*APP*; Swedish: K670N, M671L, Florida: I716V, and London: V717I) and two in presenilin 1 (*PSEN1*; M146L and L286V).

Female B6.Cg-Tg mice were crossed to male BXD mice to produce what we call AD-BXD F1 progeny (Table 1). We used four types of sires—BXD32/TyJ (RRID:IMSR_JAX:000078), BXD65/RwwJ (RRID:IMSR_JAX:007110), BXD77/RwwJ (RRID:IMSR_JAX:007121), and BXD101/RwwJ (RRID:IMSR_JAX:007144), all sourced from the UTHSC BXD colony. The terminology of these F1 hybrids is complex, but to give the reader some guidance, a 5XFAD female crossed to a male BXD65 has the charming official designation: (B6.Cg-Tg(APPSwFlLon,PSEN1*M146L*L286V)6799Vas/Mmjax x BXD65/RwwJ)F1/J.

In table 1 we highlight only the sire of the cross, because the dam is always a C57BL/6J congenic female who carries a single hemizygous copy of the 5XFAD transgenes (Fig. 1, Methods). All five of the parental strains have been deeply sequenced and all F1 hybrids therefore have completely defined genomes ^13 128^. The BXD32 strain has an unusual genetic history and due to a backcross early in its lineage has a genome of which 75% is derived from DBA/2J. The set of all four types of AD-BXD F1s (AD-BXD32 F1, AD-BXD65 F1, AD-BXD77 F1, and AD-yBXD101 F1) segregate for approximately 5.7 million SNPs and other short variants. They inherit one copy of all autosomes, Chr X, and their mitochondrial genome from the C57BL/6J congenic dam. Other autosomes and Chr Y in males are inherited from the BXD sire (Fig. 1a). At any locus F1 progeny (cases and controls) are either homozygous for the *B* reference allele or heterozygous for *B* and *D* alleles. On average half of progeny will inherit the hemizygous bigenic 5XFAD transgene located on Chr 3 (Tg cases) and half will not—the non-transgenic controls (NTg). We refer to both types of F1s as AD-BXD F1s.

Female maternal congenic mice were used so that the mitochondrial genome was constant, and to control for strain-specific differences in pre- and postnatal maternal effects, such as maternal behavior.

Male and female offspring were group housed (2–5 per cage) and maintained on a 12 hr light/dark cycle with *ad libitum* access to food and water. All mice were genotyped at weaning in-house using JAX Protocol 31769 (www.jax.org/protocol?stocknumber=008730&protocolid=31769).

All mouse experiments were carried out at University of Tennessee Health Science Center in accordance with the standards of the Association for the Assessment and Accreditation of Laboratory Animal Care (AAALAC), as well as the recommendations of the National Institutes of Health Guide for the Care and Use of Laboratory Animals. The protocol was approved by the Institutional Animal Care and Use Committee (IACUC) at the University of Tennessee Health Science Center.

### Behavioral phenotyping

Mice were phenotyped at approximately 14 months of age using a battery of physiological and behavioral assays over a two-week period, including in order of testing: body weight, grip strength, open-field testing, what-where-which testing, rotarod testing, Y-maze testing, snowflake maze testing, and finally, context fear memory acquisition (CFA) and context fear memory consolidation (CFM) testing. In this paper, we mainly report on CFA and CFM.

*Contextual fear conditioning.* At the end of the behavioral battery, and after at least 10 days of habituation to transportation and the testing room, mice were trained on a standard contextual fear conditioning (CFC) paradigm. The protocol used matches that previously used in the AD-BXD F1s^12 126^. Training consisted of a 180 s baseline period followed by four mild foot shocks (1 s, 0.9 mA), separated by 115 ± 20 s. A 40 s interval following each foot shock was defined as the post-shock interval. The percentage of time spent freezing during the final post-shock interval (PS4) was used as an index of CFA, a measure of fear learning.

Twenty-four hours later, CFM was tested by returning the mouse to the testing chamber for 10 min. The percentage of time spent freezing during this trial was used as a measure of CFM, a measure known to be dependent on hippocampal function ^129^. Videos of training and test trials were captured using FreezeFrame software (Coulbourn Instruments, PA, USA). A DeepLabCut model was trained to recognize 13 points on a mouse, following the labelling system of^130^, and DLCanalyzer^130^ was used to measure freezing events. All current behavioral data are available on GeneNetwork.org by selecting SPECIES = Mouse; *Group* = BXD NIA Alzheimer’s Study; *Type* = Traits and Cofactors. Entering the string “CFM” in the *Get Any* field will retrieve GN Record ID 5XF_10474, data on CFM for over 500 mice. Every trait can be accessed with URLs of this general form where the number in the URL is the taken from the Record ID: https://genenetwork.org/show_trait?trait_id=10474&dataset=BXD-NIA-ADPublish

### Sacrifice and perfusion

Following behavioral testing, mice were deeply anesthetized with isoflurane administered by a Somno Suite system (Kent Scientific). A 5% mixture will be used until the animal is deeply anesthetized, and the surgical procedure will not begin until the animal is no longer able to respond to pain (toe/tail pinches) and trancardially perfused with heparinized PBS for 5 min followed by PBS-4% formaldehyde to which ProHance gadolinium (Gadoteridol, Bracco, Inc.) for another 5 min. ProHance reduces the spin-lattice relaxation time (T1), enabling a shorter repetition time and greatly reducing scan times. The cranium without skin was stored for 24 hours at 4°C in 13 ml of neutral buffered formalin. The following day the mandible and muscles were trimmed away, and the crania and parts of the maxillary region including the eyes and periorbital regions were placed in 1% PBS with 1% ProHance and shipped by air from UTHSC to Duke in on ice in pressure-proof containers. Crania remained in ProHance for at least an addition two weeks prior to scanning.

### MR histology

Details of imaging methods are provided in Ref.^29^ and are outlined in **Fig. 1c-m**). Diffusion tensor imaging (DTI) was performed with a 9.4T vertical bore MRI system designed for MR histology (**Fig. 1c**).This system features a high-performance gradient (Resonance Research, Inc. BFG 88-41) providing peak gradients of 3000 mT/m. 3D volumes were acquired using a Stejskal-Tanner spin echo sequence^131^ at 25 µm isotropic resolution (**Fig. 1d**). Compressed sensing with an 8X acceleration^115^ and high angular resolution diffusion encoding (HARDI)^132^ was used with gradients (b = 3000 s/mm²) sampled along 61 axes to obtain uniform angular coverage of a unit sphere. Baseline (b0) images were generated after every 10th diffusion-weighted scan (except the last 10), resulting in a total of 66 3D volumes. During the first two years of data gathering (2021– 2022), the scanner was controlled by an Agilent Direct Drive console. The system was upgraded in 2023 to an MR Solutions console.

Automated imaging pipelines were crucial for managing the large data volumes. A script initiated on the scanner (**Fig. 1c**) triggered the scan and started data transfer to the high-performance compute cluster (**Fig. 1g**). When all the data for the first volume had been transferred to the cluster, a Fourier transform was performed along the read-out axis separating the 3D volume into many (∼800) 2D files which were then distributed across the cluster cores for parallel iterative reconstruction. At the same time, the script loaded the next set of acquisition parameters (e.g., b value, angle) and initiated the next scan. This process ran unattended for approximately 22 h facilitating high throughput with minimal operator errors.

Once the scans were completed, a pipeline built on ANTs^133^ registered the 66 volumes into a common space to correct for small displacements caused by eddy currents in the magnet, which vary between volumes. This step was critical due to the small voxel size and the strength of the gradients (up to 50 times stronger than standard clinical DTI). The 4D volume was then denoised using a machine learning algorithm^134^ and processed through DSI Studio (dsi-studio.labsolver.org), where diffusion scalar images were generated using a DTI algorithm^135^ (**Fig. 1e**).

After all scans were complete, each specimen was labeled using another automated pipeline based on ANTs^133^. A hybrid 3D volume was created for each specimen by combining the diffusion-weighted image (DWI)—that is to say, the average of all the diffusion images and the fractional anisotropy image. To minimize bias from individual specimens, a pairwise minimum deformation template (MDT) was generated using the hybrid images for all individuals of a particular genetic background type. This MDT was then registered to the Duke Mouse Brain Atlas, a stereotaxic population atlas of the 90-day male C57BL/6J mouse, which was created from high-resolution (15 µm) DTI data, microCT, and complementary light-sheet imaging ^2^. Labels were placed on the MDT, and the transformations for each specimen were inverted. The reduced common coordinate reference labels (RCCFv3)^29^ are a subset of the Allen Mouse Brain Common Coordinate Framework (CCFv3)^35^ which have been put into a framework with stereotaxic accuracy^2^.

### Empirical estimation of the precision of ROI delineations using left-right coefficients of variation

We compute CVs of independently segmented right and left volumes of each ROI—the standard deviation of the two values divided by their mean. Like standard deviations, CVs are biased downward by about 25% with a sample size of only two ^136^ but we did not apply this correction because the technical components of CVs are overestimated by biological sources of asymmetry. Note that we claim *precision* of delineation rather than *accuracy*. Accuracy would require an additional step to confirm that delineations are not only precise but that they conform to consensus boundaries established by a group of skilled neuroanatomists—a sociological enterprise that has proved to be problematic for over 150 years. But we have spot-checked consistency of MRH-driven delineations of volumes with those generated by skilled neuroscientists and stereologists (e.g., Figs. 2, 3). For example, our delineations of the dorsal lateral geniculate nucleus (LGD) of AD-BXD F1s have mean unilateral volumes of approximately 0.35 ± 0.04 SD mm^3^, whereas our previous stereological estimates of adult B6D2 F1 hybrid from celloidin-sectioned Nissl-stained material are 0.32 ± 0.03 mm^3^ with correction for shrinkage ^137^. Those of Evangelino and colleagues^87^ are about 0.26 mm^3^ for adult C57BL/6J mice as delineated in serial Nissl-stained sections without correction for shrinkage. We estimate that the LGD volume of this particular strain with correction of shrinkage is about 15% higher (0.30 ± 0.02 mm^3^, see Ref ^137^). From genetic and statistical perspectives precision and replicability are operationally more critical than accuracy in terms of detecting causes and effects of volumetric changes.

### Light Sheet Microscopy

Brains were removed from the skull, placed in buffered formalin, and shipped to LifeCanvas Technologies (Cambridge, Mass) in airtight containers. Specimens were cleared using SHIELD^138^ and labeled using eFLASH technology which integrates stochastic electrotransport ^139^ and SWITCH methods^140^). Brains were labeled using 180 µg Cell Signaling Technology rabbit beta amyloid (aβ) (RRID:AB_2798671), 120 µg EnCor Biotechnology mouse NeuN (RRID:AB_2572267), and 480 µL Millipore goat ChAT (RRID:AB_2079751). After index matching the samples were imaged using a SmartSPIM axially-swept light sheet microscope using a 3.6x objective (0.2 NA) (LifeCanvas Technologies, Cambridge, MA) using three different excitation sources (488 nm for beta amyloid; 561 nm for NeuN; and 647 nm for ChAT) producing three tiff files each of about 300 GB. Data were transmitted via Globus to a dedicated alignment server at the CIVM (**Fig. 1g**). The inevitable distortions in light sheet processed tissue and images were removed by aligning the NeuN (561 nm) volumes to the original DWI volumes of the same individuals (Fig 1h). Details of this workflow are included in Ref.^27^. The transforms generated in this alignment were applied to the other two channels—aβ and ChAT. The delineations of the ROIs were then transferred to the LSM volumes. We are able to estimate densities of aβ, and NeuN+ cells in anatomically correct stereotaxic space using a digital version of direct 3-dimensional counting, also known as the optical dissector^28,141^.

### Statistical Analyses

ANOVA was performed for each terminal ROI and for all parent ROIs (Fig. 4a), for example all ROIs contributing to the thalamus (THA). We used a workflow developed with MATLAB 2021b. The primary metrics that we evaluated were absolute ROI volumes and the fractional volume of the whole brain—the ratio of the bilateral volume of each ROI for each individual brain given the total volume of that brain. The overall ANOVA model used in most analyses (Fig. 4) evaluates sources of variance of fractional volumes (bilateral sums of ROIs) as a function of transgene status, genetic background, sex, and MRI scanner. The last factor was added because we upgraded the scanner console during the study, but we note that this factor was insignificant in all analyses. Benjamini-Hochberg (BH) corrections were used to define the *P* values at FDRs ranging from 0.05 in figure 3, to more lenient values for exploratory and comparative analyses in figure 4.

Additional stratified ANOVA tests were performed to consider the sex and strain effects separately on the most significantly affected ROIs at several BH corrected *P* values from the overall ANOVA test. For the strain-stratified ANOVAs, we used the model: fractional volume ∼ transgene status + sex + scanner + error. The AD-BXD101 cases and controls were only scanned on the newer console from MR Solutions Group (www.mrsolutions.com),and therefore does not have a scanner covariate. For the sex-stratified ANOVAs, we used the model: fractional volume ∼ transgene status + strain + scanner. Effect size measures are provided for all ANOVA tests as Cohen’s F of the transgene status and as Cohen’s D providing a pairwise comparison of key classes (cases vs controls, males vs. females).

Correlations between ROI fraction volume and either CFA or CFM were calculated using Spearman rank correlations, implemented in R (www.R-project.org). To compare the correlations in cases vs controls, we used the *cocor* R package^142^, which makes use of Fisher’s *r*-to-Z transformation which has been found to be a robust way to compare Spearman rank correlations.

Data can be accessed at the individual level at GeneNetwork.org^143^, under the group heading of *BXD NIA Alzheimer’s Studies (BXD-NIA-AD)* (see Supplementary Table READ ME). All brain volume data are available under assertion GN1068, and behavioral phenotypes used in this study are CFA (5XF_10594) and CFM (5XF_10474). All data are freely available under FAIR guidelines ^144^.

## Supporting information

Read me

